# Tumors Negate the Action of ImpL2 by Elevating Wingless

**DOI:** 10.1101/2020.09.16.299255

**Authors:** Jiae Lee, Katelyn G.-L. Ng, Kenneth M. Dombek, Young V. Kwon

## Abstract

Tumors often secrete wasting factors associated with atrophy and degeneration of host tissues. If tumors were affected by the wasting factors, mechanisms allowing tumors to evade the adverse effects of the wasting factors must exist and impairing such mechanisms may attenuate tumors. We used *Drosophila* midgut tumor models to show that tumors upregulate Wingless (Wg) to oppose the growth-impeding effects caused by the wasting factor, ImpL2 (Insulin-like growth factor binding protein (IGFBP)-related protein). Growth of Yorkie (Yki)-induced tumors is dependent on Wg while either elimination of *ImpL2* or elevation of Insulin/IGF signaling in tumors revokes this dependency. Notably, Wg augmentation could be a general mechanism for supporting the growth of tumors with elevated ImpL2 and exploited to attenuate muscle degeneration during wasting. Our study elucidates the mechanism by which tumors negate the action of ImpL2 and implies that targeting the Wnt/Wg pathway might be an efficient treatment strategy for cancers with elevated IGFBPs.

## Introduction

Reduction of tissue mass is a hallmark of the tissue wasting associated with cancers and other chronic conditions (Argiles et al., 2014; Baracos et al., 2018; Peixoto da Silva et al., 2020; Penna et al., 2014). Notably, a key feature of muscle wasting is catabolism of muscle proteins, which results in muscle mass loss (Baracos et al., 2018; Peixoto da Silva et al., 2020). Reduction in tissue mass is also observed in *Drosophila* models of cachexia-like wasting (Chatterjee and Deng, 2019; Figueroa-Clarevega and Bilder, 2015; Kreipke et al., 2017; Kwon et al., 2015; Saavedra and Perrimon, 2019). Formation of tumors in the midgut by expression of an active form of the transcription factor in the Hippo pathway, Yorkie (Yki^3S/A^) (Oh and Irvine, 2009), or transplantation of malignant disc tumors into adult flies induces atrophy of ovaries and fat body (Figueroa-Clarevega and Bilder, 2015; Kwon et al., 2015). These tumors express a high level of the secreted *Drosophila* insulin-like peptides (Dilps) antagonist ImpL2, which contributes to tissue atrophy by reducing systemic Insulin/Insulin-like growth factor (IGF) signaling (Figueroa-Clarevega and Bilder, 2015; Kreipke et al., 2017; Kwon et al., 2015). Similarly, a decrease in IGF-1 signaling in wasting muscle is well documented in mammals (Bodine et al., 2001; Costelli et al., 2006; Schiaffino et al., 2013). Diminishing Akt activity in the muscle leads to activation of the Forkhead transcription factor (FoxO) and Autophagy-related 1 (Atg1), which in turn increases protein catabolism (Fearon et al., 2012; Peixoto da Silva et al., 2020; Penna et al., 2013; Schiaffino et al., 2013). Multiple secreted factors contributing to tissue wasting during cachexia have been identified (Argiles et al., 2014; Fearon et al., 2013; Peixoto da Silva et al., 2020). In particular, the transforming growth factor β family members, Activins, induce muscle protein catabolism in part by inhibiting Akt signaling (Chen et al., 2014; Han et al., 2013; Zhou et al., 2010). Given that the growth of cancers accompanies an increase in mass via activation of various anabolic processes, the organismal state under cachexia is expected to be unfavorable for cancer growth; however, it has been shown that cancers grow in a variety of cancer cachexia models (Das et al., 2011; Gallot et al., 2014; Kir et al., 2014; Zhou et al., 2010). Conceptually, if cancers were to respond to the secreted wasting factors, these factors would oppose cancer growth. Thus, cancers must have a mechanism to overcome the adverse effects induced by the wasting factors to ensure their growth during cachexia. Nevertheless, it is unclear whether these wasting factors could oppose cancer growth during cachexia and how cancers evade the potentially growth-impeding effects induced by the wasting factors to uphold their growth.

In *Drosophila*, binding of Dilps to Insulin-like receptor (InR) initiates the Insulin/IGF pathway by turning on Phosphoinositide 3-kinases (PI3K), which leads to activation of Akt (Akt1 in *Drosophila*) (Nassel et al., 2015). Akt1 activation promotes growth by inhibiting the *Drosophila* Forkhead transcription factor Foxo, which is a growth suppressor, and activating Target of rapamycin (Tor), which is a conserved regulator of cell size and organ growth (Nassel et al., 2015). In turn, Tor inhibits Thor (4E-BP in humans) to enhance translational initiation and activates Ribosomal protein S6 kinase (S6k) to increase ribosome biogenesis (Nassel et al., 2015). Additionally, Tor suppresses autophagy by inhibiting Atg1 (Chang and Neufeld, 2010). Thus, maintaining Insulin/IGF signaling is crucial for supporting growth of tissues as well as an organism. In contrast, attenuation of Insulin/IGF signaling is associated with tissue wasting in *Drosophila* (Dionne et al., 2006; Figueroa-Clarevega and Bilder, 2015; Kwon et al., 2015). Recent studies have demonstrated that ImpL2 is a tumor-derived wasting factor, which induces a reduction in systemic Insulin/IGF signaling (Figueroa-Clarevega and Bilder, 2015; Kwon et al., 2015). One puzzling observation is that Yki^3S/A^-induced midgut tumors (*yki*^*3S/A*^ tumors) and transplanted malignant disc tumors can grow regardless of the dramatic increase in ImpL2 expression in these tumors. Considering the fundamental role of Insulin/IGF signaling in growth, these tumors must have a mechanism to evade the growth-impeding effect induced by ImpL2. It is not known how these tumors maintain Insulin/IGF signaling even though ImpL2 is highly upregulated in these tumors.

In this study, we employ *Drosophila* midgut tumor models to address whether tumors are also subjected to the ramifications of ImpL2 elevation and how these tumors overcome the ImpL2-mediated adverse effect on their growth. Interestingly, it has been shown that eye disc tumors use Wingless (Wg) signaling to promote tumor progression in the condition of insulin resistance induced by a high-sugar diet (Hirabayashi et al., 2013). Given that tumor-derived ImpL2 causes a reduction in systemic Insulin/IGF signaling, reminiscent of insulin resistance (Figueroa-Clarevega and Bilder, 2015; Kwon et al., 2015), we address the role of Wg in the growth of midgut tumors. Our observations demonstrate that Wg is necessary for increasing Insulin/IGF signaling in *yki*^*3S/A*^ tumors. Notably, Wg is indispensable for the growth of *yki*^*3S/A*^ tumors. This indispensability can be abolished by either depleting *ImpL2* or increasing Insulin/IGF signaling in *yki*^*3S/A*^ tumors. Finally, we show that the Wg-mediated antagonism of the ImpL2 action could be a general principle for supporting the growth of a subtype of midgut tumors with elevated ImpL2 expression and be exploited to alleviate muscle degeneration during cachexia-like wasting.

## Results

### Wg is essential for the growth of *yki*^*3S/A*^ tumors only in the presence of ImpL2

To address the role of Wg signaling in the growth of *yki*^*3S/A*^ tumors, we first assessed whether Wg was expressed in tumors which were generated by expressing *yki*^*3S/A*^ with *esg-GAL4, UAS-GFP, tub-GAL80*^*ts*^ (referred to as *esg*^*ts*^, hereafter; see methods). Previous studies have demonstrated that Wg is expressed in the visceral muscle and the intestinal epithelial compartment (Cordero et al., 2012; Lin et al., 2008). Wg expressed from the visceral muscles is essential for homeostatic intestinal stem cell (ISC) self-renewal (Lin et al., 2008). In contrast, Wg expressed in enteroblasts (EBs) during tissue damage plays a crucial role in epithelial regeneration (Cordero et al., 2012). Comparison of *wg* mRNA levels in control and *yki*^*3S/A*^ tumor midguts revealed that *wg* mRNA expression was significantly elevated in *yki*^*3S/A*^ tumors (Figure 1A). To discern which compartment of *yki*^*3S/A*^ tumor midguts expressed Wg, we stained the midguts with an anti-Wg antibody. Wg signals were significantly elevated in *esg*^*+*^ cells upon expression of *yki*^*3S/A*^ with *esg*^*ts*^ (Figure 1B). In contrast, Wg signals in the visceral muscle remained unchanged in *yki*^*3S/A*^ tumor midguts (Figure S1). Altogether, these results indicate that Wg is elevated in *yki*^*3S/A*^ cells.

**Figure 1.**
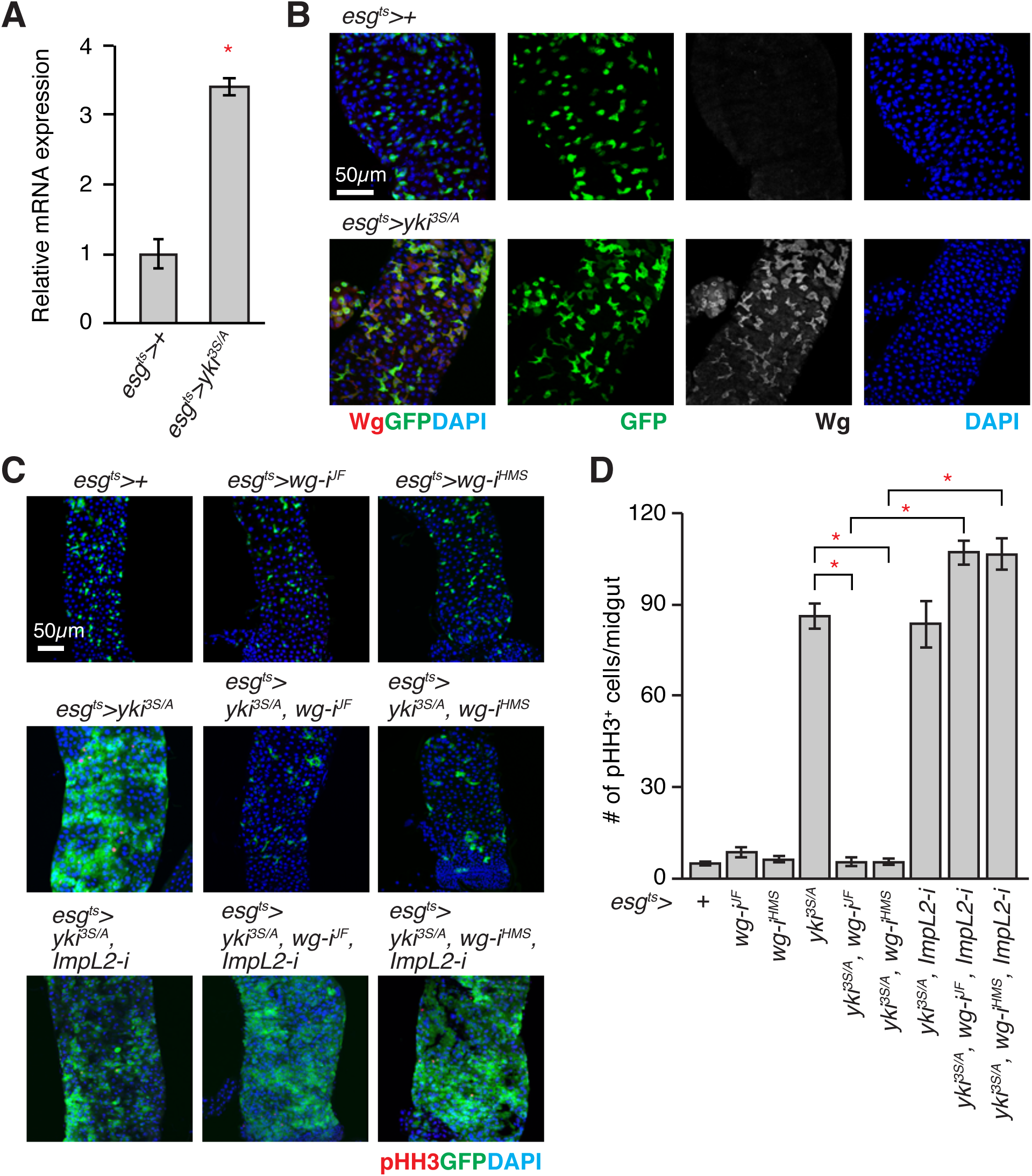
*Wg* is indispensable for the growth of *yki*^*3S/A*^ tumor in the presence of *ImpL2*. (A) Expression of *wg* mRNA in midguts. Relative abundance of *wg* transcript in *esg*^*ts*^*>+* or *esg*^*ts*^*>yki*^*3S/A*^ midguts was determined by qRT-PCR after 3 days of transgene expression. (B) Immunostaining of Wg in posterior midguts. Transgenes were induced for 3 days. The cells manipulated by *esg*^*ts*^ are marked by GFP (green), Wg staining is shown in red, and nuclei are stained with DAPI (blue) in merged images. Scale bar, 50 *µ*m. (C) Representative images of posterior midguts. Transgenes were expressed for 5 days with *esg*^*ts*^. Phospho-Histone H3 (PHH3) staining is shown in red. (D) Quantification of pHH3^+^ cells per midgut. RNAi lines: *wg-i*^*JF*^, *JF01257*; *wg-i*^*HMS*^, *HMS00794*; *ImpL2-i, 15009R-3*. N=20 (*esg*^*ts*^*>+*), N=11 (*esg*^*ts*^*>wg-i*^*JF*^), N=11 (*esg*^*ts*^*>wg-i*^*HMS*^), N=22 (*esg*^*ts*^*>yki*^*3S/A*^), N=12 (*esg*^*ts*^*>yki*^*3S/A*^, *wg-i*^*JF*^), N=10 (*esg*^*ts*^*>yki*^*3S/A*^, *wg-i*^*HMS*^), N=12 (*esg*^*ts*^*>yki*^*3S/A*^, *wg-i*^*JF*^, *ImpL2-i*), N=12 (*esg*^*ts*^*>yki*^*3S/A*^, *wg-i*^*HMS*^, *ImpL2-i*), N=22 (*esg*^*ts*^*>yki*^*3S/A*^, *ImpL2-i*) biological replicates. Mean±SEMs are shown. \**P*<0.01, two-tailed unpaired Student’s *t*-test compared with control (*esg*^*ts*^*>+*) unless indicated by bracket. See also Figure S1.

To test the role of Wg in *yki*^*3S/A*^ tumor growth, we depleted *wg* in *yki*^*3S/A*^ cells using two independent *wg* RNA interference (RNAi) lines: *JF01257* and *HMS00794* (Chen et al., 2016; Lee et al., 2014). Knockdown of *wg* in ISCs and EBs had no effect on homeostatic ISC division (Figures 1C and 1D). In contrast, expression of *wg* RNAi in *yki*^*3S/A*^ tumors with *esg*^*ts*^ significantly reduced cell proliferation, resulting in a complete suppression of tumor growth (Figures 1C and 1D).

ImpL2 antagonizes Dilps, which leads to a reduction in Insulin/IGF signaling, with an exception in a small subset of neurons in the larval brain (Amoyel et al., 2016; Bader et al., 2013; Honegger et al., 2008; Nie et al., 2019; Okamoto et al., 2013; Sarraf-Zadeh et al., 2013). Given the suggested role of Wg in promoting the progression of eye disc tumors under insulin resistance induced by a high-sugar diet (Hirabayashi et al., 2013), we hypothesized that *yki*^*3S/A*^ tumor-derived Wg might negate the adverse effect caused by ImpL2 elevation to support tumor growth. If this is correct, the tumor-growth defect caused by *wg* depletion should be rescued by *ImpL2* depletion in *yki*^*3S/A*^ tumors. Previously, we showed that *ImpL2* was dispensable for the growth of *yki*^*3S/A*^ tumors (Kwon et al., 2015). Consistently, *ImpL2* depletion didn’t altered the proliferation of *yki*^*3S/A*^ cells (Figure 1C). Of significance, expression of *ImpL2* RNAi with *esg*^*ts*^ completely rescued the defect in *yki*^*3S/A*^ cell proliferation caused by *wg* knockdown, leading to the formation of fully-grown tumors (Figures 1C and 1D). Altogether, these results demonstrate that Wg is crucial for the growth of *yki*^*3S/A*^ tumors only when *ImpL2* is present.

### Augmentation of Insulin-Akt signaling is sufficient to rescue the tumor-growth defect caused by *wg* depletion

Given the complete rescue of the tumor-growth defect by *ImpL2* depletion, we hypothesized that Wg supports *yki*^*3S/A*^ tumor growth by mainly negating the action of ImpL2. It has been shown that Wg expressed in disc tumors increases Insulin/IGF signaling by increasing the expression of *Insulin-like peptide receptor* (*InR*) (Hirabayashi et al., 2013). Thus, we explored whether Wg could affect Insulin/IGF signaling in intestinal ISCs and EBs. While we were testing the effect of *wg* knockdown, we noticed that expression of *wg* RNAi with *esg*^*ts*^ significantly reduced the size and the number of *esg*^*+*^ cells (Figures 2A and 2B). If these phenotypes were mediated by a reduction in Insulin/IGF signaling, augmenting Insulin/IGF signaling in ISCs and EBs should reverse the phenotypes. Indeed, expression of a constitutively active *Akt1* (*myr-Akt1*) with *esg*^*ts*^ rescued the phenotypes caused by *wg* depletion in ISCs and EBs (Figures 2A and 2B). Next, we tested whether increasing Wg was sufficient to induce an elevation in Insulin/IGF signaling in ISCs and EBs. Ectopic expression of Wg with *esg*^*ts*^ caused an increase in phospho-Akt1 signals in *esg*^*+*^ cells (Figure 2C). Altogether, these results suggest that Wg produced from *esg*^*+*^ cells plays an important role in regulating Insulin/IGF signaling in ISCs and EBs, and ectopic Wg expression is sufficient to increase Akt1 phosphorylation in ISCs and EBs.

**Figure 2.**
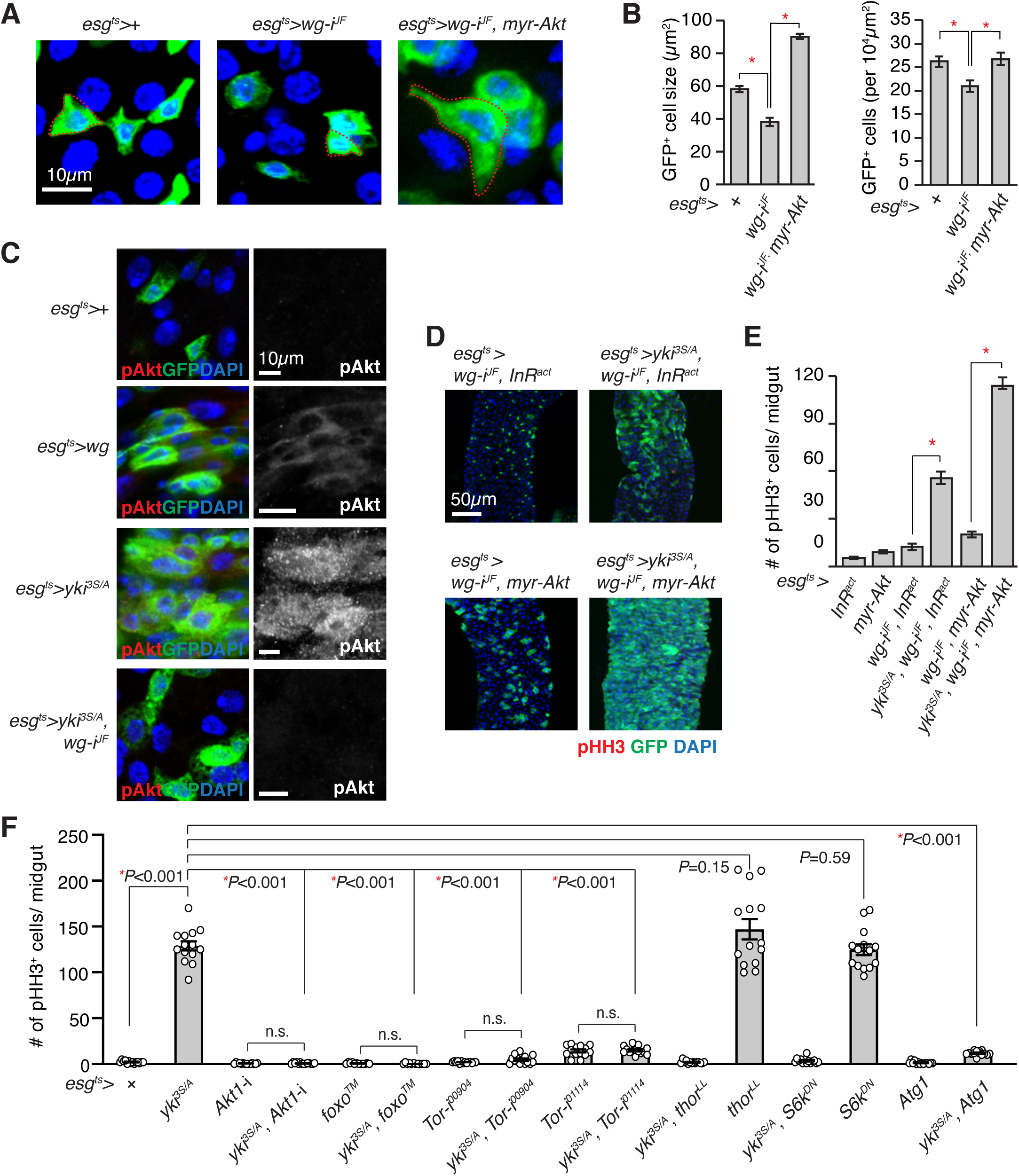
Activation of Insulin/IGF signaling rescues the *yki*^*3S/A*^ tumor growth defect caused by *wg* depletion. (A) Representative images of *esg*^*+*^ cells. Red dotted line indicates cell boundary. (B) Quantification of cell size and number. The size and number of *esg*^*+*^ cells in 100×100 *µ*m^2^ were quantified. (C) Phospho-Akt (pAkt) immunostaining. Transgenes were expressed for 5 days. Scale bar, 10 *µ*m. (D) Representative images of posterior midguts after 5 days of transgene expression with *esg*^*ts*^. GFP (green) marks *esg*^*+*^ cells, and pHH3 signals are shown in red. Scale bar, 50 *µ*m. (E) Quantification of pHH3^+^ cells in midguts. N=9 (*esg*^*ts*^*>InR*^*act*^), N=20 (*esg*^*ts*^*>myr-Akt1*), N=13 (*esg*^*ts*^*>wg-i*^*JF*^, *InR*^*act*^), N=19 (*esg*^*ts*^*>yki*^*3S/A*^, *wg-i*^*JF*^, *InR*^*act*^), N=17 (*esg*^*ts*^*>wg-i*^*JF*^, *myr-Akt*)1, N=11 (*esg*^*ts*^*>yki*^*3S/A*^, *wg-i*^*JF*^, *myr-Akt1*) biological replicates. (F) Quantification of pHH3^+^ cells per midgut after 5 days of transgene expression. For (B), (E) and (F), mean±SEMs are shown. \**P*<0.01, two-tailed unpaired Student’s *t*-test between two genotypes indicated by bracket. See also Figure S2.

Previously, it was shown that phospho-Akt1 levels were increased in *yki*^*3S/A*^ tumors relative to control midguts while phospho-Akt1 levels in the host muscle, ovaries, and heads were significantly decreased in flies bearing *yki*^*3S/A*^ tumors in the midgut (Kwon et al., 2015). In accordance with the previous observations, phospho-Akt1 signals were increased in *yki*^*3S/A*^ cells compared to control cells (Figure 2C). Notably, we found that *wg* depletion in *yki*^*3S/A*^ cells significantly reduced phospho-Akt1 signals, suggesting that the increase in phospho-Akt1 signals in *yki*^*3S/A*^ cells was dependent on Wg (Figure 2C). If the role of Wg in supporting *yki*^*3S/A*^ tumor growth is to oppose the effect caused by ImpL2 elevation via increasing Insulin/IGF signaling, augmenting Insulin/IGF signaling in *yki*^*3S/A*^ cells should be sufficient to rescue the growth defect caused by *wg* depletion. Of significance, ectopic expression of either an active form of *InR* (*InR*^*act*^) or *myr-Akt1* rescued the defect in *yki*^*3S/A*^ tumor growth caused by *wg* depletion (Figures 2D and 2E). These results demonstrate that Wg is necessary for increasing Insulin/IGF signaling in *yki*^*3S/A*^ tumors, which is important for negating the action of ImpL2.

### Activation of Foxo or Atg1 attenuates *yki*^*act*^ tumor growth

Since our observations indicate that Wg supports *yki*^*3S/A*^ tumor growth by increasing Insulin/IGF signaling, we decided to investigate which branch of the Insulin/IGF pathway is essential for *yki*^*3S/A*^ tumor growth. In control *esg*^*+*^ cells, *Akt1* depletion caused a reduction in homeostatic ISC division (Figures 2F and S2). Similarly, *Akt1* depletion in *yki*^*3S/A*^ cells led to a complete suppression of *yki*^*3S/A*^ tumor growth (Figures 2F and S2). Interestingly, expression of *Akt1* RNAi in combination of *yki*^*3S/A*^ with *esg*^*ts*^ almost completely eliminated *esg*^*+*^ cells while expression of *Akt1* RNAi alone with *esg*^*ts*^ caused only a slight reduction in the number of *esg*^*+*^ cells (Figure S2). Ectopic expression of a mutant foxo (foxo™) which cannot be phosphorylated by Akt1 (Hwangbo et al., 2004) or depletion of *Tor* significantly decreased the division of *yki*^*3S/A*^ cells (Figures 2F and S2A). To elucidate the Tor downstream mediator that is essential for *yki*^*3S/A*^ tumor growth, we manipulated three well-characterized Tor downstream players. Expression of either a mutant *thor* (*thor*^*LL*^) which cannot be inhibited by Tor due to the mutations at the mTOR phosphorylation sites (Miron et al., 2001) or a dominant negative S6k (S6k^DN^) (Barcelo and Stewart, 2002) in *yki*^*3S/A*^ cells didn’t significantly alter the division of *yki*^*3S/A*^ cells (Figure 2F and S2A). Of significance, ectopic expression of Atg1 in *yki*^*3S/A*^ cells almost completely abolished *yki*^*3S/A*^ tumor growth (Figure 2F and S2A). These results suggest that attenuation of the Foxo and Atg1 signaling branches in the Insulin/IGF pathway is critical for supporting *yki*^*3S/A*^ tumor growth.

### *Wg* is specifically upregulated in tumors with elevated *ImpL2* expression

Our observations suggest that *yki*^*3S/A*^ tumor growth is dependent on Wg upregulation, which might restrain Foxo signaling and Atg1 signaling by antagonizing the action of ImpL2. Interestingly, a recent study demonstrated that Atg1 could inhibit Yki by direct phosphorylation (Tyra et al., 2020), raising the possibility that elevation of Wg could be a mechanism specifically applicable to *yki*^*3S/A*^ tumors. Considering the fundamental role of Yki in controlling the growth of tumors (Snigdha et al., 2019; Zheng and Pan, 2019), Wg upregulation might be a general mechanism to support the growth of tumors, especially with elevated ImpL2 expression. Thus, we decided to test whether a similar mechanism exists to support the growth of other types of midgut tumors.

In addition to *yki*^*3S/A*^ midgut tumors, we found that midgut tumors driven by expression of a combination of *Ras*^*V12*^ and dominant negative *Notch* (*N*^*DN*^) with *esg*^*ts*^ (*esg*^*ts*^/*UAS-N*^*DN*^; *UAS-Ras*^*V12*^/+) induced systemic organ wasting, which was manifested by the bloating syndrome phenotype, fat body degeneration, and ovary atrophy (Figure 3A). In contrast, tumors driven by expression of *N*^*DN*^ alone, a gain of function allele of *Raf* (*Raf*^*gof*^), or *Unpaired* (*Upd1*) and *Signal-transducer and activator of transcription protein at 92E* (*Stat92E*) didn’t induce discernable wasting phenotypes (Figure 3A). Consistent with the proposed role of ImpL2 in systemic organ wasting, *ImpL2* mRNA expression was increased greater than 80-fold in *Ras*^*V12*^, *N*^*DN*^ tumors compared to control midguts while it remained unaltered in *Raf*^*gof*^ and *Upd1, Stat92E* tumors (Figure 3B). Note that a moderate but significant increase in *ImpL2* mRNA levels was also observed with *N*^*DN*^ tumors (Figure 3B). If upregulation of Wg is a general mechanism by which tumors evade the adverse effects caused by ImpL2 elevation, Wg expression would be expected to be increased in tumors with an elevated ImpL2 expression. Accordingly, we detected a strong correlation between *wg* and *ImpL2* mRNA levels (R^2^=0.9732 and r=0.9864); *wg* mRNA expression was increased specifically in *Ras*^*V12*^, *N*^*DN*^ and *N*^*DN*^ tumors while it was unaltered in the other tumors (Figure 3B). Furthermore, Wg protein signals were cell-autonomously increased in *Ras*^*V12*^, *N*^*DN*^ and *N*^*DN*^ cells (Figure 3C).

**Figure 3.**
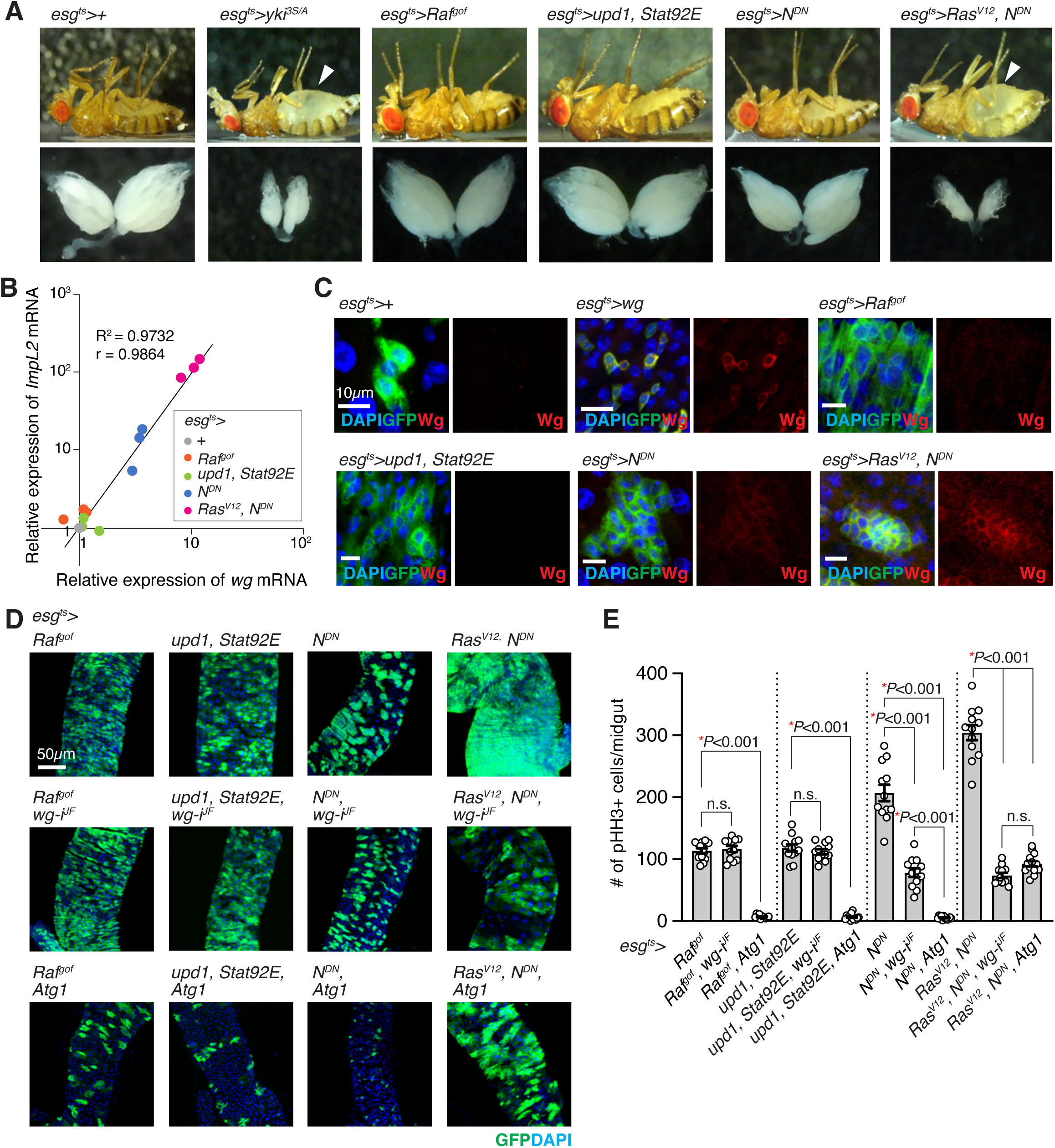
Wg is specifically required for the growth of midgut tumors with elevated ImpL2 expression. (A) Representative images of fly and ovary. Arrowheads indicate abdominal bloating. *esg*^*ts*^>*Ras*^*V12*^, *N*^*DN*^ flies were incubated at 29°C for 4 days, and control (*esg*^*ts*^>+) and other flies were incubated for 6 days to induce transgene expression. (B) Correlative plot of relative *ImpL2* and *wg* mRNA levels. mRNA expression values of *wg* and *ImpL2* in the midguts with indicated genotypes were measured by qRT-PCR and then normalized to those values in the control midguts (*esg*^*ts*^*>*+). Relative *wg* and *ImpL2* mRNA levels from three independent experiments are shown in x-axis and y-axis, respectively. The coefficient of determination (R^2^) is 0.9732, and the Pearson correlation coefficient (r) is 0.9864 (P<0.0001). (C) Wg immunostaining in the midguts. Scale bars, 10 *µ*m. (D) Images of posterior midguts. Scale bar, 50 *µ*m. (E) Quantification of pHH3^+^ cells per midgut. Mean±SEMs are shown with individual data points. \**P*<0.01, two-tailed unpaired Student’s *t*-test between two groups indicated with bracket. All transgenes were induced with *esg*^*ts*^ for 6 days except for ‘*Ras*^*V12*^, *N*^*DN*^*’*, ‘*Ras*^*V12*^, *N*^*DN*^, *wg-i*^*JF*^*’*, ‘*Ras*^*V12*^, *N*^*DN*^, *Atg1*’, which were induced for 4 days.

### Wg is specifically required for the growth of tumors with *ImpL2* elevation

To assess the importance of Wg on the growth of these tumors, we depleted *wg* by expressing *wg* RNAi with *esg*^*ts*^. Notably, the growth of *Raf*^*gof*^ and *Upd1, Stat92E* tumors was unaltered by *wg* depletion (Figures 3D and 3E). In contrast, expression of *wg* RNAi with *esg*^*ts*^ significantly suppressed the growth of both *Ras*^*V12*^, *N*^*DN*^ and *N*^*DN*^ tumors (Figures 3D and 3E); a more prominent suppression was observed with *Ras*^*V12*^, *N*^*DN*^ tumors, which expressed significantly higher levels of *ImpL2* mRNA (Figure 3C). Since we could generate a few different midgut tumors in the absence of direct manipulation of Yki, we sought to address the effect of Atg1 activation on the growth of these tumors. Interestingly, ectopic expression of Atg1 with *esg*^*ts*^ significantly suppressed the growth of all the tumors (Figures 3D and 3E). Taken together, these results suggest that Wg elevation is specifically required for the growth of the midgut tumors with elevated ImpL2 expression while attenuation of Atg1 appears to be a general requirement for the growth of midgut tumors.

### Ectopic Wg expression in the muscle increases Insulin/IGF signaling and rescues muscle degeneration in the flies harboring *yki*^*3S/A*^ midgut tumors

*yki*^*3S/A*^ tumors in the midgut induce muscle degeneration, which is dependent on tumor-derived ImpL2 (Kwon et al., 2015). Given the observation that Wg expression with *esg*^*ts*^ could increase Akt1 phosphorylation in *esg*^*+*^ cells (Figure 2C), we sought to address whether ectopic expression of Wg in the muscle could rescue muscle degeneration induced by *yki*^*3S/A*^ tumors in the midgut. To express Wg in the muscle while simultaneously inducing *yki*^*3S/A*^ tumors in the midgut, we established a LexA::GAD-based temperature-sensitive inducible system (*StanEx*^*SX-4*^ (Kockel et al., 2016), *LexAop-mCD8::GFP, tub-GAL80*^*ts*^, hereafter referred as *esg-LexA::GAD*^*ts*^, see methods) and a transgenic line harboring *LexAOP-yki*^*3S/A*^. As a result, we were able to generate *yki*^*3S/A*^ tumors (*esg-LexA::GAD*^*ts*^*/+; LexAOP-yki*^*3S/A*^*/+*) independent of a GAL4/UAS system (Figure S4A). To manipulate gene expression in the muscle, we used *Mhc*.*F3-580-GAL4*, which has been shown to express GAL4 mainly in the adult indirect flight muscle (Gajewski and Schulz, 2010).

It has been previously shown that *yki*^*3S/A*^ midgut tumors caused a disparity in Insulin/IGF signaling between *yki*^*3S/A*^ tumors and host tissues in an ImpL2-dependent manner: Phospho-Akt signals were significantly reduced in host tissues while increased in *yki*^*3S/A*^ tumors (Kwon et al., 2015). Wg expression using *Mhc*.*F3-580-GAL4* while inducing *yki*^*3S/A*^ tumors in the midgut (*esg-LexA::GAD*^*ts*^*/Mhc*.*F3-580-GAL4; LexAOP-yki*^*3S/A*^*/UAS-wg*) increased phospho-Akt signals specifically in the muscle; phospho-Akt signals in ovaries, fat body, and the neighboring muscle compartment were unaltered (Figures 4A and S3). Notably, we found that Wg expression with *Mhc*.*F3-580-GAL4* significantly increased mRNA levels of not only *InR* but also *Akt1* and *chico* in the muscle (Figure 4B), which might contribute to the increase in Dilp sensitivity in the muscle (Hirabayashi et al., 2013; Zhang et al., 2011). Expression of Wg in the muscle had no effect on the growth of *yki*^*3S/A*^ tumors in the midgut (Figure S4). Thus, our observations suggest that ectopic expression of Wg is sufficient to increase Insulin/IGF signaling in the muscle.

**Figure 4.**
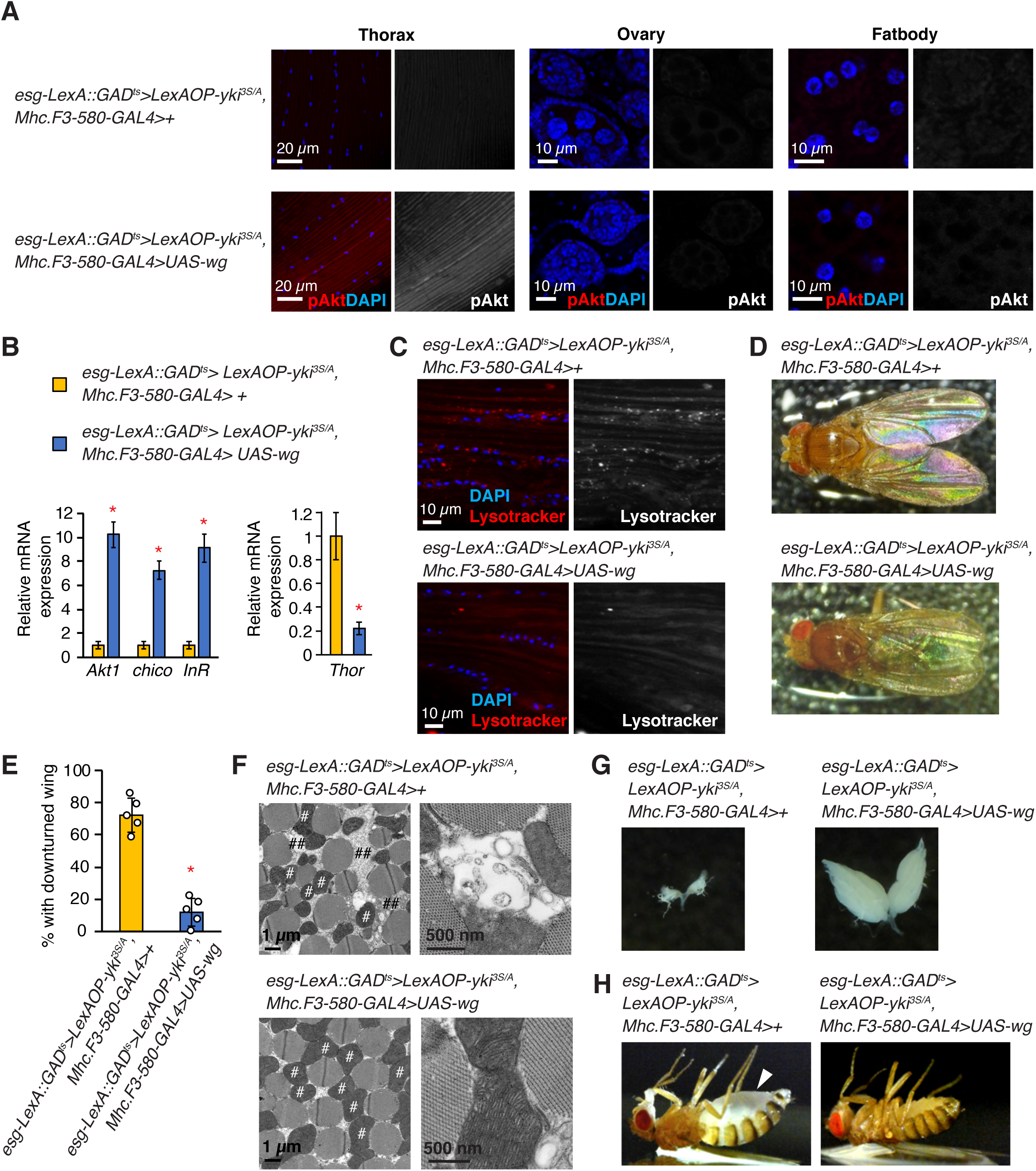
Ectopic Wg expression rescues muscle degeneration caused by *yki*^*3S/A*^ tumors in the midgut. (A) Phospho-Akt staining. Thoraces from male flies and ovaries and fat body from female flies were used. Phospho-Akt signals are shown in red in merge, and nuclei are stained with DAPI (blue). (B) Relative mRNA expression in the thorax. Mean±SEMs are shown. \**P*<0.01, two-tailed unpaired Student’s *t*-test. (C) Lysotracker staining in thorax. Lysotracker signals are shown in red in merged images, and nuclei are stained with DAPI (blue). Scale bar, 10 *µ*m. (D) Dorsal view of male flies, (E) Penetrance of downturned wing phenotype. N=69 (*esg-LexA::GAD*^*ts*^*/Mhc*.*F3-580-GAL4; LexAOP-yki*^*3S/A*^*/+*), N=71 (*LexA::GAD*^*ts*^*/Mhc*.*F3-580-GAL4; LexAOP-yki*^*3S/A*^*/UAS-wg*), male flies were used for 5 independent experiments. Mean±SEMs are shown with individual data points. \**P*<0.01, two-tailed unpaired Student’s *t*-test. (F) Electron microscopic images of the transverse section of indirect flight muscles. # denotes mitochondria; ## denotes low electron-dense sector between myofibers. (G) Representative images of ovary. (H) Representative images of flies. Arrowhead indicates abdominal bloating. All transgenes were induced for 8 days. See also Figures S3 – S5.

Reduction in Insulin/IGF signaling can lead to an activation of both Foxo and Atg1 in host tissues. Although Foxo signaling has been shown to be upregulated in the muscle of the flies bearing *yki*^*3S/A*^ tumors (Kwon et al., 2015), it is not known whether Atg1 signaling is also activated in the host tissues. Strong lysotracker signals were detected in the host tissues prepared from *yki*^*3S/A*^ tumor-bearing flies (Figure S5). Depleting *ImpL2* in *yki*^*3S/A*^ tumors was sufficient to suppress the accumulation of lysotracker signals in the host tissues (Figure S5), suggesting that induction of autophagy in the host tissues was dependent on ImpL2 derived from *yki*^*3S/A*^ tumors. Of significance, Wg expression in the muscle reduced lysotracker signals in the muscle of *yki*^*3S/A*^ tumor-bearing flies, an indicative of attenuation of Atg1 signaling (Figure 4C). Additionally, Wg expression in the muscle of *yki*^*3S/A*^ tumors-bearing flies significantly reduced *thor* mRNA levels (Figure 4B). Our results indicate that ectopic expression of Wg in muscle could decrease both Foxo and Atg1 activities.

Strikingly, increasing Wg in the muscle was sufficient to inhibit muscle degeneration in *yki*^*3S/A*^ tumors-bearing flies, manifested by down-turned wing phenotype and muscle mitochondrial fragmentation (Figures 4D and 4E). Wg expression in the muscle also rescued ovary atrophy and the bloating syndrome phenotype induced by *yki*^*3S/A*^ tumors (Figures 4G and 4H). Thus, these results demonstrate that augmenting Wg in the muscle can rescue not only muscle degeneration but also a few wasting phenotypes observed outside of muscle.

## Discussion

In this study, we demonstrate that Wg upregulation is the mechanism by which *yki*^*3S/A*^ tumors evade the growth-impeding effects induced by ImpL2. Our observations indicate that the main role of Wg in supporting the growth of *yki*^*3S/A*^ tumors is to increase Insulin/IGF signaling in a tumor-autonomous manner. Therefore, without Wg, *yki*^*3S/A*^ tumors are influenced by the action of ImpL2, which can be attenuated by either depleting *ImpL2* or augmenting Insulin/IGF signaling in *yki*^*3S/A*^ tumors (Figures 1C, 1D, 2D, and 2E). Of significance, we observed a strong correlation between Wg and ImpL2 expression levels in several types of midgut tumors; Wg expression was increased in only those tumors with elevated *ImpL2* expression (Figures 3B and 3C). Notably, *wg* depletion specifically affected the growth of the midgut tumors with elevated ImpL2 expression (Figures 3D and 3E), suggesting that upregulation of Wg might be a general mechanism for supporting the growth of the midgut tumors with elevated ImpL2 expression.

We have previously shown that tumor-derived ImpL2 plays a key role in inducing the disparity in Insulin/IGF signaling between *yki*^*3S/A*^ tumors and host tissues by reducing systemic Insulin/IGF signaling (Kwon et al., 2015). Based on our findings described in this study, we propose that Wg upregulation in *yki*^*3S/A*^ tumors is also a crucial factor for inducing the disparity in Insulin/IGF signaling. This disparity increases Foxo and Atg1 activities in the host tissues even though the role of Foxo and Atg1 in cachexia-like wasting still needs to be addressed in *Drosophila*. Nevertheless, our observations suggest that Wg-mediated upregulation of Insulin/IGF signaling might be important for restraining both Foxo and Atg1 activities in *yki*^*3S/A*^ tumors, which is critical for supporting tumor growth (Figure 2F and S2A).

We demonstrate that ectopic expression of Wg in the muscle rescues muscle degeneration in *yki*^*3S/A*^ tumors-bearing flies. Wg expression in the muscle increased Akt1 phosphorylation specifically in the muscle compartment, but not in ovaries or fat body (Figure 4A). Additionally, expression of Wg with *esg*^*ts*^ increased Akt1 phosphorylation in *esg*^*+*^ cells, not in the neighboring enterocytes (ECs) (Figure 2C). This cell autonomous action of Wg might account for the rescue of muscle degeneration by Wg expression (Figures 4D and 4F). It is intriguing to note that augmenting Wg in the muscle also rescued ovary degeneration and the bloating syndrome phenotype (Figures 4G and 4H). We speculate that potential interorgan communication between degenerating muscle and other tissues might be the basis of the Wg-mediated rescue of the wasting phenotypes outside the muscle.

Taken together, our study provides insights into the mechanism by which tumors negate the action of the tumor-derived wasting factor ImpL2. Interestingly, the mammalian ImpL2-like proteins IGFBPs are upregulated in multiple types of cancers, and multiple IGFBPs are shown to function as tumor suppressors in mammals (Chan et al., 2018; Wajapeyee et al., 2008; Wang et al., 2015; Wu et al., 2015; Zumkeller, 2001). Considering that the conserved role of Wnt/Wg signaling in increasing IGF signaling (Abiola et al., 2009; Inoki et al., 2006; Palsgaard et al., 2012), it would be intriguing to investigate whether a similar relationship between Wnt and IGFBPs exists in human cancers.

## Acknowledgements

We thank Dr. N. Perrimon for the fly stocks and the valuable comments, Dr. Y. Kim for the help with fly crosses, and Dr. E. Parker for electron microscopy. This work was supported by Mallinckrodt grant to Y.V.K from the Edward Mallinckrodt, Jr. Foundation and the Core Grant for Vision Research (NEI P30EY001730).

## Author Contributions

J.L. and Y.V.K. designed experiments, analyzed data, and wrote the manuscript. J.L. and K.G.-L.N. performed experiments. K.M.D. generated a DNA construct.

## KEY RESOURCES TABLE

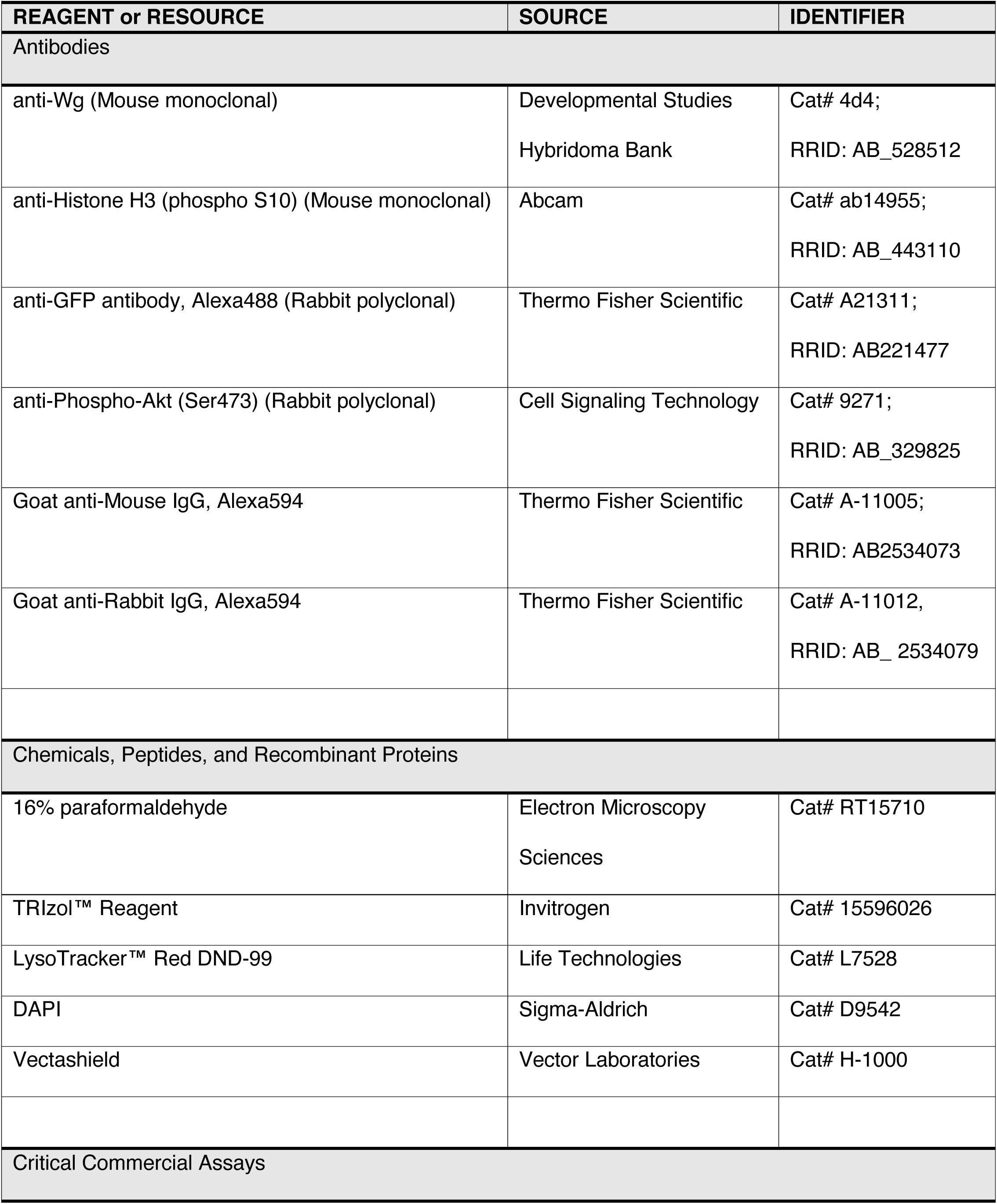
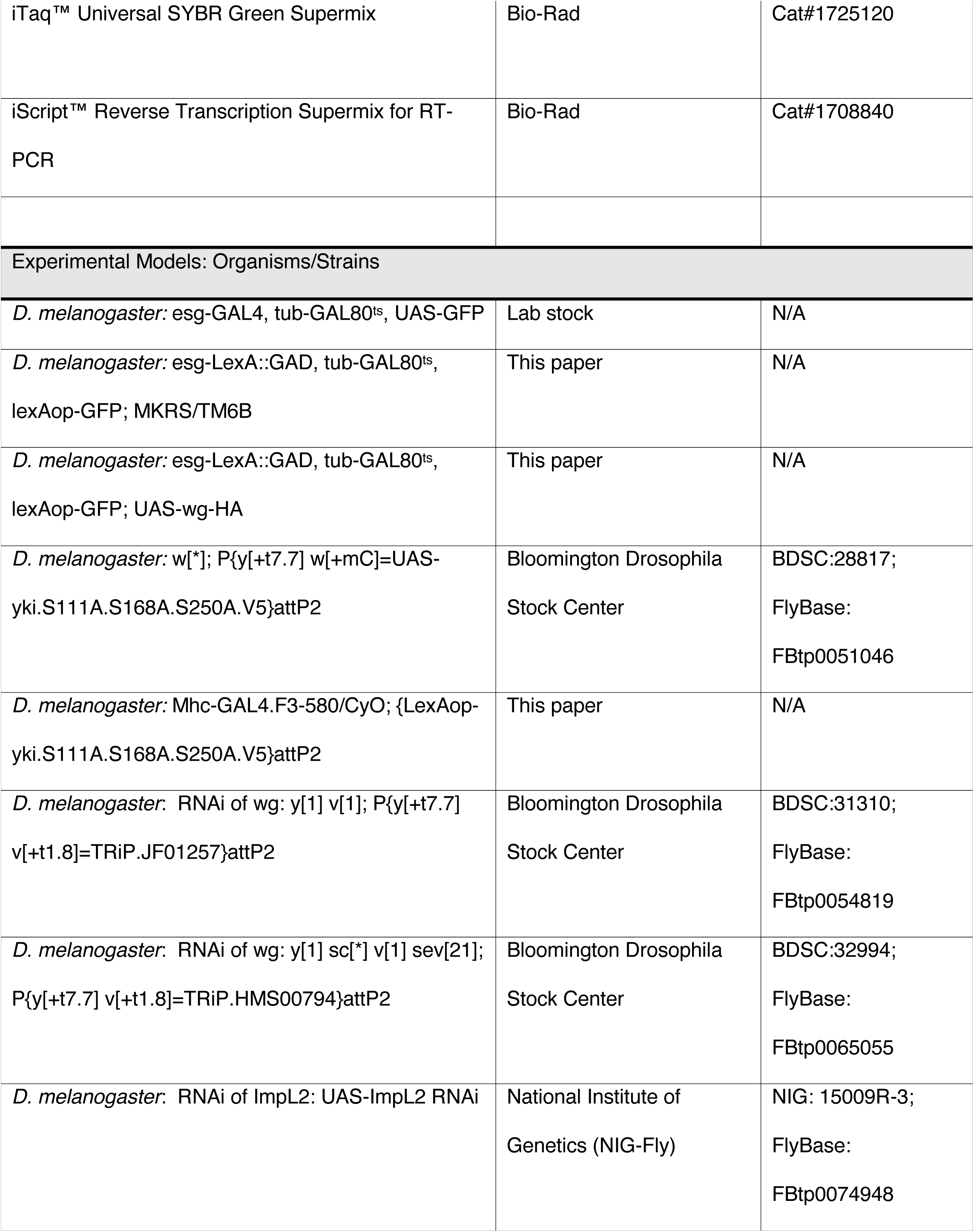
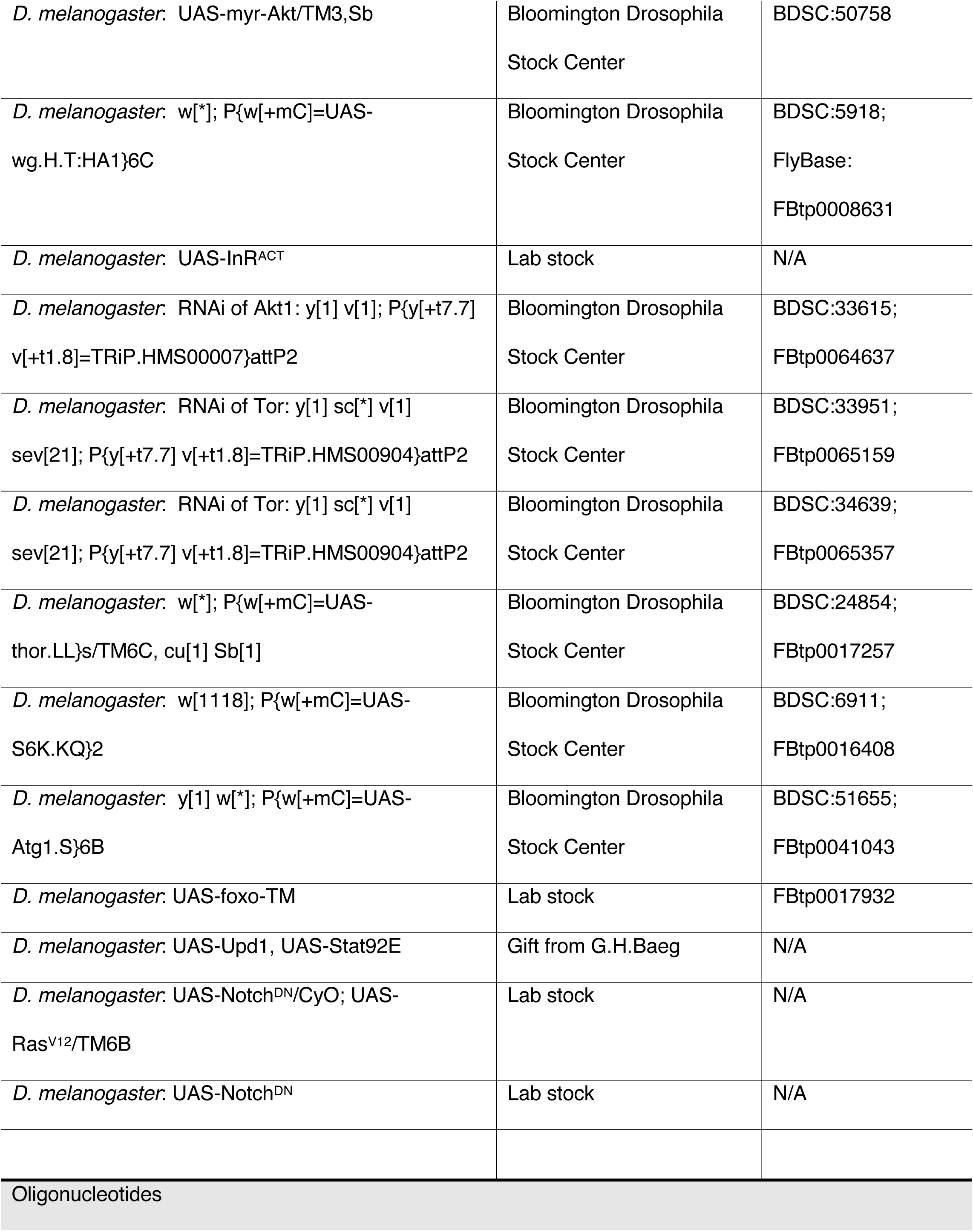
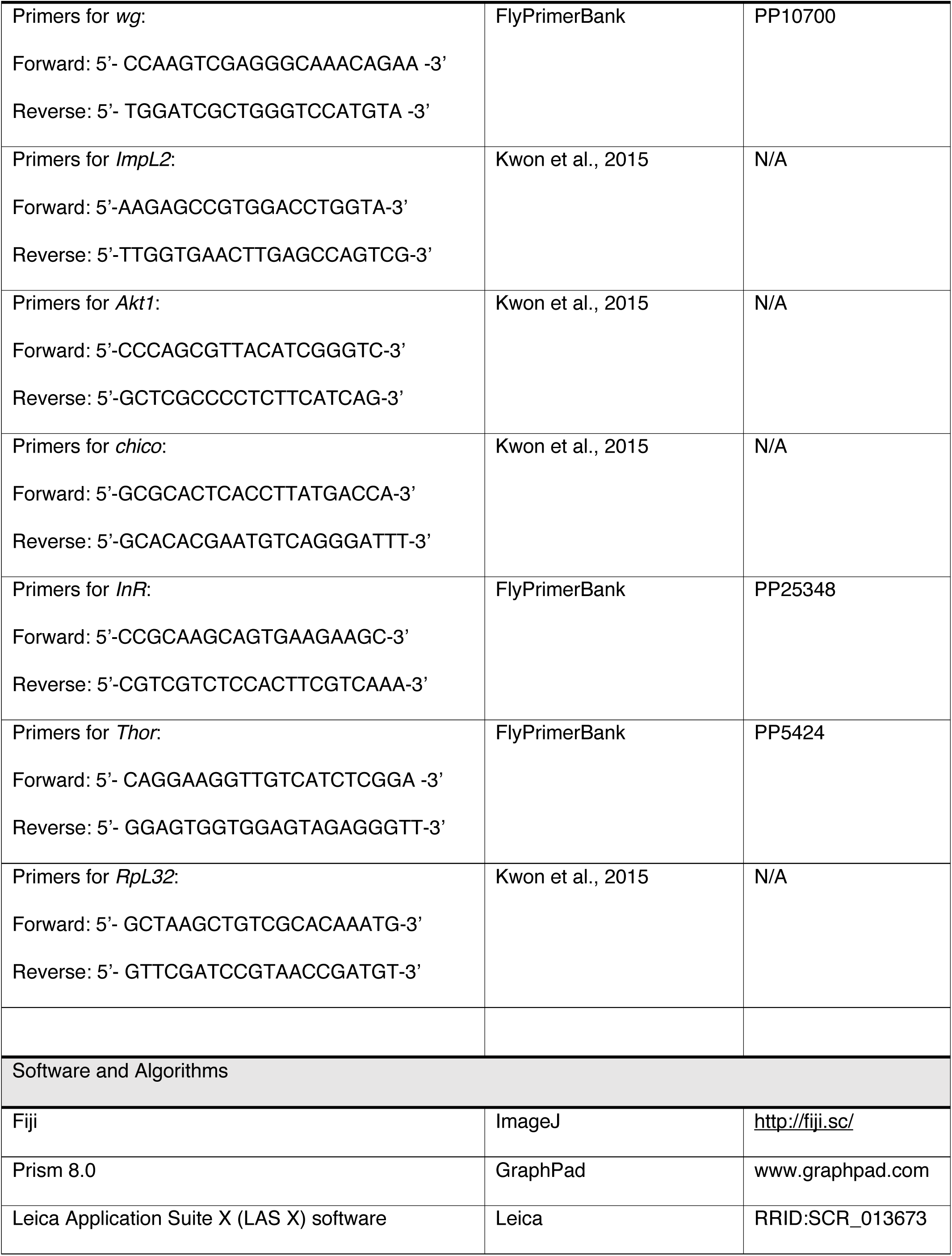

### Star⋆Methods

#### CONTACT FOR REAGENT AND RESOURCE SHARING

Further information and requests for reagents may be directed to and will be fulfilled by the corresponding author Young V. Kwon.

#### EXPERIMENTAL MODEL AND SUBJECT DETAILS

Several lines of the fruit fly *Drosophila melanogaster* were used in this study and are listed in the Key Resources Table. Fly crosses were set up in vials containing standard molasses-agar medium, kept at room temperature for 3 days, and then transferred to 18°C to restrict the expression of GAL4-induced transgenes throughout the development. Adult progenies were collected and incubated at 29°C for 3-to 8-days prior to dissection to induce the transgenes. During incubation at 29**°**C, flies were transferred onto fresh food every 2 days. Female flies (<16 day old) were used for all experiments except for those employed in thorax, where male flies were used instead.

To manipulate intestinal stem cells (ISCs) and enteroblasts (EBs), we used *esg-GAL4, tub-GAL80*^*ts*^, *UAS-GFP* (referred as *esg*^*ts*^) and *StenEx*^*SX-4*^, *tub-GAL80*^*ts*^, *LexAop-mCD8::GFP*, (referred as *esg-lexA*::*HG*^*ts*^*). StenEx*^*SX-4*^ (BDSC#66659) was recombined with *tub-GAL80*^*ts*^ (BDSC#7108) and *LexAop2-mCD8::GFP* (BDSC#32205) to generate *esg-lexA*::*HG*^*ts*^. Other *Drosophila* lines and their sources are listed in the Key Resources Table.

## METHOD DETAILS

### Generation of *LexAOP-yki*^*3S/A*^ Line

We obtained the *yki* ^*S111A*.*S168A*.*S250A*^ (referred as *yki*^*3S/A*^) coding sequence from Dr. Irvine at Rutgers University (reference 2009 oncogene). The *yki*^*3S/A*^ sequence was amplified by PCR using the following primers: Forward: 5’-ctcgagATGTTAACGACGATGTCAGCCAG-3’, Reverse: 5’-tctagattaATTAATTTTATACCATTCCAAATCGTCAGG-3’. The PCR product was subcloned into pJFRC19-13XLexAOP-IVS-myr::GFP vector (Addgene#26224) to generate pJFRC19-13XLexAOP-*yki*^*3S/A*^. The resulting construct was targeted into the attP2 site through germline transformation.

### Quantitative RT-PCR

Total RNA from adult female midguts and male thoraces was isolated with TRIzol (Invitrogen, Cat# 15596026). RNA (1*µ*g) was used to produce cDNA with iScript™ Reverse Transcription Supermix (Bio-Rad, Cat#1725120). The cDNA was subjected to quantitative real-time PCR with iTaq™ Universal SYBR Green Supermix (Bio-Rad, Cat#1708840) and CFX-96 (Bio-Rad). *RpL32* was used for normalization. The fold change in RNA expression compared to the control was calculated and plotted for relative mRNA expression. Primers used for qRT-PCR are described in the Key Resource Table.

### Antibody staining and immunofluorescence microscopy

To remove food from the midgut, flies were fed on 4% sucrose for approximately 4 hours prior to dissection. We prepared midguts, ovaries and fat body from Female flies and thoraxes from male flies. Tissues dissected in PBS were fixed in 4% paraformaldehyde (Electron Microscopy Sciences) for 20 minutes and then washed 3 times for 5 minutes each with PBST (PBS supplemented with 0.2% Triton X-100). For permeabilization and blocking, tissue samples were incubated in blocking buffer (PBST supplemented with 5% normal goat serum) for 1 hour at room temperature. Then, tissue samples were incubated with primary antibody in blocking buffer overnight at 4**°**C. The tissue samples were washed three times with PBST for 5 minutes each, and then incubated with secondary antibody for 2–3 hours at room temperature. Stained tissues were washed 3 times with PBST and mounted with Vectashield (Vector Laboratories, Cat# H-1000). Fluorescence micrographs were acquired with Leica SP8 laser scanning confocal microscope with 40x/1.25 oil objective lens. Fiji software was used for further adjustment and assembly of the acquired images.

### Lysotracker staining

For lysotracker staining, thoraces of male flies were dissected in four pieces: cut once in sagittal section and once in transverse section, in PBS. Female flies were used for ovary staining and female abdominal cuticle was used for fat body staining. Freshly dissected tissues were incubated for 5 min in 50 nM LysoTracker Red DND-99 (Life Technologies, Cat# L7528) in PBS, rinsed quickly three times in PBS, and then fixed in 4% PFA in PBS for 5 minutes in room temperature. The samples were briefly incubated in PBST for permeabilization, stained with DAPI, rinsed three times with PBST, and mounted in Vectashield. Muscle fibers were carefully dissociated and removed from the thoracic cuticle to spread flat when mounting.

### Quantification of phospho-Histone H3 (pHH3)-positive cells

To determine the number of cells undergoing mitotic division, midguts were dissected and stained with anti-pHH3 antibody (Abcam, Cat# ab14955). Number of pHH3-positive nuclei were counted from the entire midgut.

### Measurement of cell size

Outlines of individual cells from confocal images acquired with 40x/1.25 oil objective lens was traced, and area was measured with Fiji software.

### Electron microscopy

Thoraces from male flies were dissected and fixed overnight in in 4% glutaraldehyde in 0.1 M sodium cacodylate buffer, pH 7.2. The samples were washed in buffer, postfixed in 1% osmium tetroxide for 90 min, rinsed, stained in 1% uranyl acetate, dehydrated in ethanol solutions, and embedded in epoxy resin (Epon Araldite). Serial sections (80 nm) were aligned and viewed on JEOL-1230 transmission electron microscope with AMT XR80 camera.

## QUANTIFICATION AND STATISTICAL ANALYSIS

All the midgut images presented and used for quantification were obtained from the posterior R5 region of female flies, except for the pHH3-positive nuclei quantification, which was done from the entire midguts. Statistical analyses were performed using Microsoft Excel and GraphPad Prism 8. All *P* values were determined by two-tailed Student’s *t*-test with unequal variances. Statistical significance was depicted by asterisks in the figures: \**P*< 0.01. Sample sizes were chosen empirically based on the observed effects and indicated in the figure legends.

## DATA AND SOFTWARE AVAILABILITY

The authors declare that the data supporting the findings of this study are available within the paper and its supplementary information files. Additional datasets are available from the corresponding author upon reasonable request.

## References

Abiola, M., Favier, M., Christodoulou-Vafeiadou, E., Pichard, A.L., Martelly, I., and Guillet-Deniau, I. (2009). Activation of Wnt/beta-catenin signaling increases insulin sensitivity through a reciprocal regulation of Wnt10b and SREBP-1c in skeletal muscle cells. PLoS One 4, e8509.

Amoyel, M., Hillion, K.H., Margolis, S.R., and Bach, E.A. (2016). Somatic stem cell differentiation is regulated by PI3K/Tor signaling in response to local cues. Development 143, 3914–3925.

Argiles, J.M., Busquets, S., Stemmler, B., and Lopez-Soriano, F.J. (2014). Cancer cachexia: understanding the molecular basis. Nat Rev Cancer 14, 754–762.

Bader, R., Sarraf-Zadeh, L., Peters, M., Moderau, N., Stocker, H., Kohler, K., Pankratz, M.J., and Hafen, E. (2013). The IGFBP7 homolog Imp-L2 promotes insulin signaling in distinct neurons of the Drosophila brain. J Cell Sci 126, 2571–2576.

Baracos, V.E., Martin, L., Korc, M., Guttridge, D.C., and Fearon, K.C.H. (2018). Cancer-associated cachexia. Nat Rev Dis Primers 4, 17105.

Barcelo, H., and Stewart, M.J. (2002). Altering Drosophila S6 kinase activity is consistent with a role for S6 kinase in growth. Genesis 34, 83–85.

Bodine, S.C., Stitt, T.N., Gonzalez, M., Kline, W.O., Stover, G.L., Bauerlein, R., Zlotchenko, E., Scrimgeour, A., Lawrence, J.C., Glass, D.J., et al. (2001). Akt/mTOR pathway is a crucial regulator of skeletal muscle hypertrophy and can prevent muscle atrophy in vivo. Nat Cell Biol 3, 1014–1019.

Chan, Y.X., Alfonso, H., Paul Chubb, S.A., Ho, K.K.Y., Gerard Fegan, P., Hankey, G.J., Golledge, J., Flicker, L., and Yeap, B.B. (2018). Higher IGFBP3 is associated with increased incidence of colorectal cancer in older men independently of IGF1. Clin Endocrinol (Oxf) 88, 333–340.

Chang, Y.Y., and Neufeld, T.P. (2010). Autophagy takes flight in Drosophila. FEBS Lett 584, 1342–1349.

Chatterjee, D., and Deng, W.M. (2019). Drosophila Model in Cancer: An Introduction. Adv Exp Med Biol 1167, 1–14.

Chen, J.L., Walton, K.L., Winbanks, C.E., Murphy, K.T., Thomson, R.E., Makanji, Y., Qian, H., Lynch, G.S., Harrison, C.A., and Gregorevic, P. (2014). Elevated expression of activins promotes muscle wasting and cachexia. Faseb J 28, 1711–1723.

Chen, Z., Zhu, J.Y., Fu, Y., Richman, A., and Han, Z. (2016). Wnt4 is required for ostia development in the Drosophila heart. Dev Biol 413, 188–198.

Cordero, J.B., Stefanatos, R.K., Scopelliti, A., Vidal, M., and Sansom, O.J. (2012). Inducible progenitor-derived Wingless regulates adult midgut regeneration in Drosophila. Embo J 31, 3901–3917.

Costelli, P., Muscaritoli, M., Bossola, M., Penna, F., Reffo, P., Bonetto, A., Busquets, S., Bonelli, G., Lopez-Soriano, F.J., Doglietto, G.B., et al. (2006). IGF-1 is downregulated in experimental cancer cachexia. American Journal of Physiology - Regulatory, Integrative and Comparative Physiology 291, R674–R683.

Das, S.K., Eder, S., Schauer, S., Diwoky, C., Temmel, H., Guertl, B., Gorkiewicz, G., Tamilarasan, K.P., Kumari, P., Trauner, M., et al. (2011). Adipose triglyceride lipase contributes to cancer-associated cachexia. Science 333, 233–238.

Dionne, M.S., Pham, L.N., Shirasu-Hiza, M., and Schneider, D.S. (2006). Akt and FOXO dysregulation contribute to infection-induced wasting in Drosophila. Curr Biol 16, 1977–1985.

Fearon, K., Arends, J., and Baracos, V. (2013). Understanding the mechanisms and treatment options in cancer cachexia. Nat Rev Clin Oncol 10, 90–99.

Fearon, K.C., Glass, D.J., and Guttridge, D.C. (2012). Cancer cachexia: mediators, signaling, and metabolic pathways. Cell Metab 16, 153–166.

Figueroa-Clarevega, A., and Bilder, D. (2015). Malignant Drosophila tumors interrupt insulin signaling to induce cachexia-like wasting. Dev Cell 33, 47–55.

Gajewski, K.M., and Schulz, R.A. (2010). CF2 represses Actin 88F gene expression and maintains filament balance during indirect flight muscle development in Drosophila. PLoS One 5, e10713.

Gallot, Y.S., Durieux, A.C., Castells, J., Desgeorges, M.M., Vernus, B., Plantureux, L., Remond, D., Jahnke, V.E., Lefai, E., Dardevet, D., et al. (2014). Myostatin gene inactivation prevents skeletal muscle wasting in cancer. Cancer Res 74, 7344–7356.

Han, H.Q., Zhou, X., Mitch, W.E., and Goldberg, A.L. (2013). Myostatin/activin pathway antagonism: molecular basis and therapeutic potential. Int J Biochem Cell Biol 45, 2333–2347.

Hirabayashi, S., Baranski, T.J., and Cagan, R.L. (2013). Transformed Drosophila cells evade diet-mediated insulin resistance through wingless signaling. Cell 154, 664–675.

Honegger, B., Galic, M., Kohler, K., Wittwer, F., Brogiolo, W., Hafen, E., and Stocker, H. (2008). Imp-L2, a putative homolog of vertebrate IGF-binding protein 7, counteracts insulin signaling in Drosophila and is essential for starvation resistance. J Biol 7, 10.

Hwangbo, D.S., Gershman, B., Tu, M.P., Palmer, M., and Tatar, M. (2004). Drosophila dFOXO controls lifespan and regulates insulin signalling in brain and fat body. Nature 429, 562–566.

Inoki, K., Ouyang, H., Zhu, T., Lindvall, C., Wang, Y., Zhang, X., Yang, Q., Bennett, C., Harada, Y., Stankunas, K., et al. (2006). TSC2 integrates Wnt and energy signals via a coordinated phosphorylation by AMPK and GSK3 to regulate cell growth. Cell 126, 955–968.

Kir, S., White, J.P., Kleiner, S., Kazak, L., Cohen, P., Baracos, V.E., and Spiegelman, B.M. (2014). Tumour-derived PTH-related protein triggers adipose tissue browning and cancer cachexia. Nature.

Kockel, L., Huq, L.M., Ayyar, A., Herold, E., MacAlpine, E., Logan, M., Savvides, C., Kim, G.E., Chen, J., Clark, T., et al. (2016). A Drosophila LexA Enhancer-Trap Resource for Developmental Biology and Neuroendocrine Research. G3 (Bethesda) 6, 3017–3026.

Kreipke, R.E., Kwon, Y.V., Shcherbata, H.R., and Ruohola-Baker, H. (2017). Drosophila melanogaster as a Model of Muscle Degeneration Disorders. Curr Top Dev Biol 121, 83–109.

Kwon, Y., Song, W., Droujinine, I.A., Hu, Y., Asara, J.M., and Perrimon, N. (2015). Systemic organ wasting induced by localized expression of the secreted insulin/IGF antagonist ImpL2. Dev Cell 33, 36–46.

Lee, J.H., Bassel-Duby, R., and Olson, E.N. (2014). Heart- and muscle-derived signaling system dependent on MED13 and Wingless controls obesity in Drosophila. Proc Natl Acad Sci U S A 111, 9491–9496.

Lin, G., Xu, N., and Xi, R. (2008). Paracrine Wingless signalling controls self-renewal of Drosophila intestinal stem cells. Nature 455, 1119–1123.

Miron, M., Verdu, J., Lachance, P.E., Birnbaum, M.J., Lasko, P.F., and Sonenberg, N. (2001). The translational inhibitor 4E-BP is an effector of PI(3)K/Akt signalling and cell growth in Drosophila. Nat Cell Biol 3, 596–601.

Nassel, D.R., Liu, Y., and Luo, J. (2015). Insulin/IGF signaling and its regulation in Drosophila. Gen Comp Endocrinol 221, 255–266.

Nie, Y., Yu, S., Li, Q., Nirala, N.K., Amcheslavsky, A., Edwards, Y.J.K., Shum, P.W., Jiang, Z., Wang, W., Zhang, B., et al. (2019). Oncogenic Pathways and Loss of the Rab11 GTPase Synergize To Alter Metabolism in Drosophila. Genetics 212, 1227–1239.

Oh, H., and Irvine, K.D. (2009). In vivo analysis of Yorkie phosphorylation sites. Oncogene 28, 1916–1927.

Okamoto, N., Nakamori, R., Murai, T., Yamauchi, Y., Masuda, A., and Nishimura, T. (2013). A secreted decoy of InR antagonizes insulin/IGF signaling to restrict body growth in Drosophila. Genes Dev 27, 87–97.

Palsgaard, J., Emanuelli, B., Winnay, J.N., Sumara, G., Karsenty, G., and Kahn, C.R. (2012). Cross-talk between insulin and Wnt signaling in preadipocytes: role of Wnt co-receptor low density lipoprotein receptor-related protein-5 (LRP5). J Biol Chem 287, 12016–12026.

Peixoto da Silva, S., Santos, J.M.O., Costa, E.S.M.P., Gil da Costa, R.M., and Medeiros, R. (2020). Cancer cachexia and its pathophysiology: links with sarcopenia, anorexia and asthenia. J Cachexia Sarcopenia Muscle 11, 619–635.

Penna, F., Baccino, F.M., and Costelli, P. (2014). Coming back: autophagy in cachexia. Curr Opin Clin Nutr Metab Care 17, 241–246.

Penna, F., Costamagna, D., Pin, F., Camperi, A., Fanzani, A., Chiarpotto, E.M., Cavallini, G., Bonelli, G., Baccino, F.M., and Costelli, P. (2013). Autophagic degradation contributes to muscle wasting in cancer cachexia. Am J Pathol 182, 1367–1378.

Saavedra, P., and Perrimon, N. (2019). Drosophila as a Model for Tumor-Induced Organ Wasting. Adv Exp Med Biol 1167, 191–205.

Sarraf-Zadeh, L., Christen, S., Sauer, U., Cognigni, P., Miguel-Aliaga, I., Stocker, H., Kohler, K., and Hafen, E. (2013). Local requirement of the Drosophila insulin binding protein imp-L2 in coordinating developmental progression with nutritional conditions. Dev Biol 381, 97–106.

Schiaffino, S., Dyar, K.A., Ciciliot, S., Blaauw, B., and Sandri, M. (2013). Mechanisms regulating skeletal muscle growth and atrophy. Febs J 280, 4294–4314.

Snigdha, K., Gangwani, K.S., Lapalikar, G.V., Singh, A., and Kango-Singh, M. (2019). Hippo Signaling in Cancer: Lessons From Drosophila Models. Front Cell Dev Biol 7, 85.

Tyra, L.K., Nandi, N., Tracy, C., and Kramer, H. (2020). Yorkie Growth-Promoting Activity Is Limited by Atg1-Mediated Phosphorylation. Dev Cell 52, 605–616 e607.

Wajapeyee, N., Serra, R.W., Zhu, X., Mahalingam, M., and Green, M.R. (2008). Oncogenic BRAF induces senescence and apoptosis through pathways mediated by the secreted protein IGFBP7. Cell 132, 363–374.

Wang, J., Ding, N., Li, Y., Cheng, H., Wang, D., Yang, Q., Deng, Y., Yang, Y., Li, Y., Ruan, X., et al. (2015). Insulin-like growth factor binding protein 5 (IGFBP5) functions as a tumor suppressor in human melanoma cells. Oncotarget 6, 20636–20649.

Wu, K., Zhou, M., Wu, Q.X., Yuan, S.X., Wang, D.X., Jin, J.L., Huang, J., Yang, J.Q., Sun, W.J., Wan, L.H., et al. (2015). The role of IGFBP-5 in mediating the anti-proliferation effect of tetrandrine in human colon cancer cells. Int J Oncol 46, 1205–1213.

Zhang, W., Thompson, B.J., Hietakangas, V., and Cohen, S.M. (2011). MAPK/ERK signaling regulates insulin sensitivity to control glucose metabolism in Drosophila. PLoS Genet 7, e1002429.

Zheng, Y., and Pan, D. (2019). The Hippo Signaling Pathway in Development and Disease. Dev Cell 50, 264–282.

Zhou, X., Wang, J.L., Lu, J., Song, Y., Kwak, K.S., Jiao, Q., Rosenfeld, R., Chen, Q., Boone, T., Simonet, W.S., et al. (2010). Reversal of cancer cachexia and muscle wasting by ActRIIB antagonism leads to prolonged survival. Cell 142, 531–543.

Zumkeller, W. (2001). IGFs and IGFBPs: surrogate markers for diagnosis and surveillance of tumour growth? Mol Pathol 54, 285–288.

