## Supplementary Information for "Tumors Negate the Action of ImpL2 by Elevating Wingless"

**Supplemental Information (Lee et al.)**

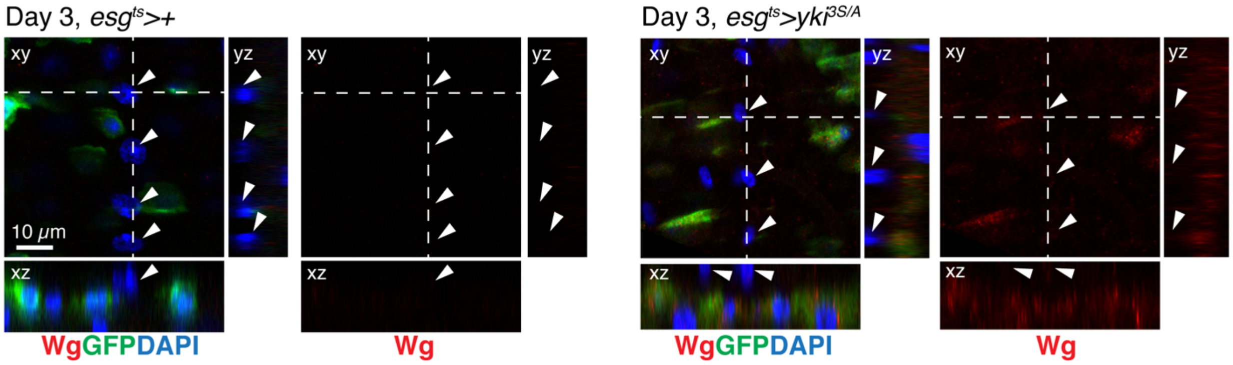


**Figure S1. *yki$^{3S/A}$* tumors do not increase Wg in visceral muscles.**

Immunostaining of Wg in posterior midguts, shown with orthogonal views. Transgenes were induced for 3 days with *esg$^{ts}$*. *esg$^{+}$* cells are marked by GFP (green), Wg staining is shown in red, and nuclei are stained with DAPI (blue) in merged images. Arrowheads point to nuclei of visceral muscle. Scale bar, 10 μm.

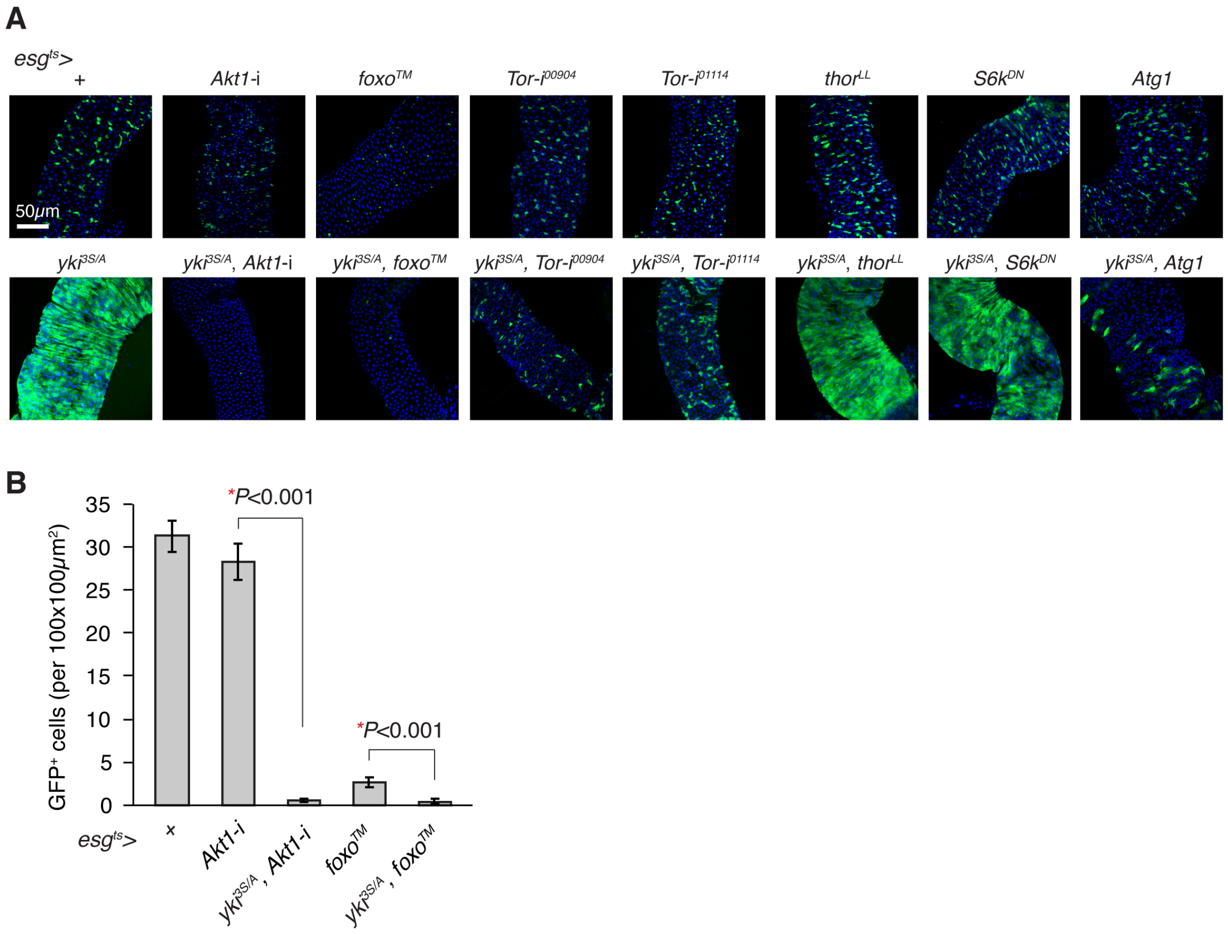


**Figure S2. Activation of Foxo or Atg1 attenuates *yki*$^{3S/A}$ tumor growth.**

(A) Representative images of posterior midguts. Transgenes were induced for 5 days with *esg*$^{ts}$. GFP (green) marks *esg*$^{+}$ cells, and nuclei are stained with DAPI (blue). RNAi lines: *Akt-i*, *HMS00007*; *Tor-i*$^{00904}$, *HMS00904*; *Tor-i*$^{01114}$, *HMS01114*. Scale bar, 50 $\mu$m.

(B) Quantification of GFP$^{+}$ cell number. The number of *esg*$^{+}$ cells in a representative 100 $\mu$mx100 $\mu$m area in each posterior midgut were quantified. N=10 (*esg*$^{ts}$>+), N=11 (*esg*$^{ts}$>*Akt-i*), N=10 (*esg*$^{ts}$>*yki*$^{3S/A}$, *Akt-i*), N=10 (*esg*$^{ts}$>*foxo*$^{TM}$), N=11 (*esg*$^{ts}$>*yki*$^{3S/A}$, *foxo*$^{TM}$). Mean±SEMs are shown. *$P$<0.01, two-tailed unpaired Student's *t*-test between two genotypes indicated by bracket.

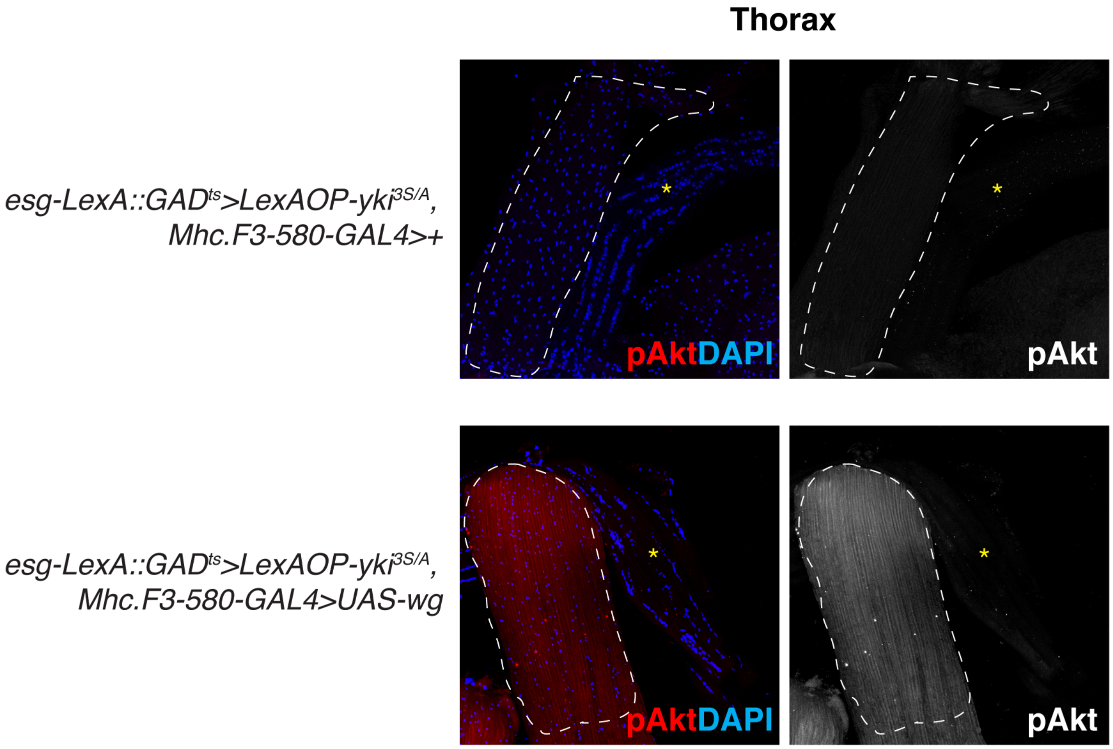


**Figure S3. Expression of Wg with *Mhc.F3-580-GAL4* increases Akt phosphorylation in the indirect flight muscle.**

*esg-LexA::GAD$^{ts}$/Mhc.F3-580-GAL4; LexAOP-yki$^{3S/A}$/wg* thorax shows elevated phospho-Akt staining in the indirect flight muscle (white dotted line), but not in the neighboring muscle compartment (asterisks). Phospho-Akt staining is shown in red, and nuclei are stained with DAPI (blue).

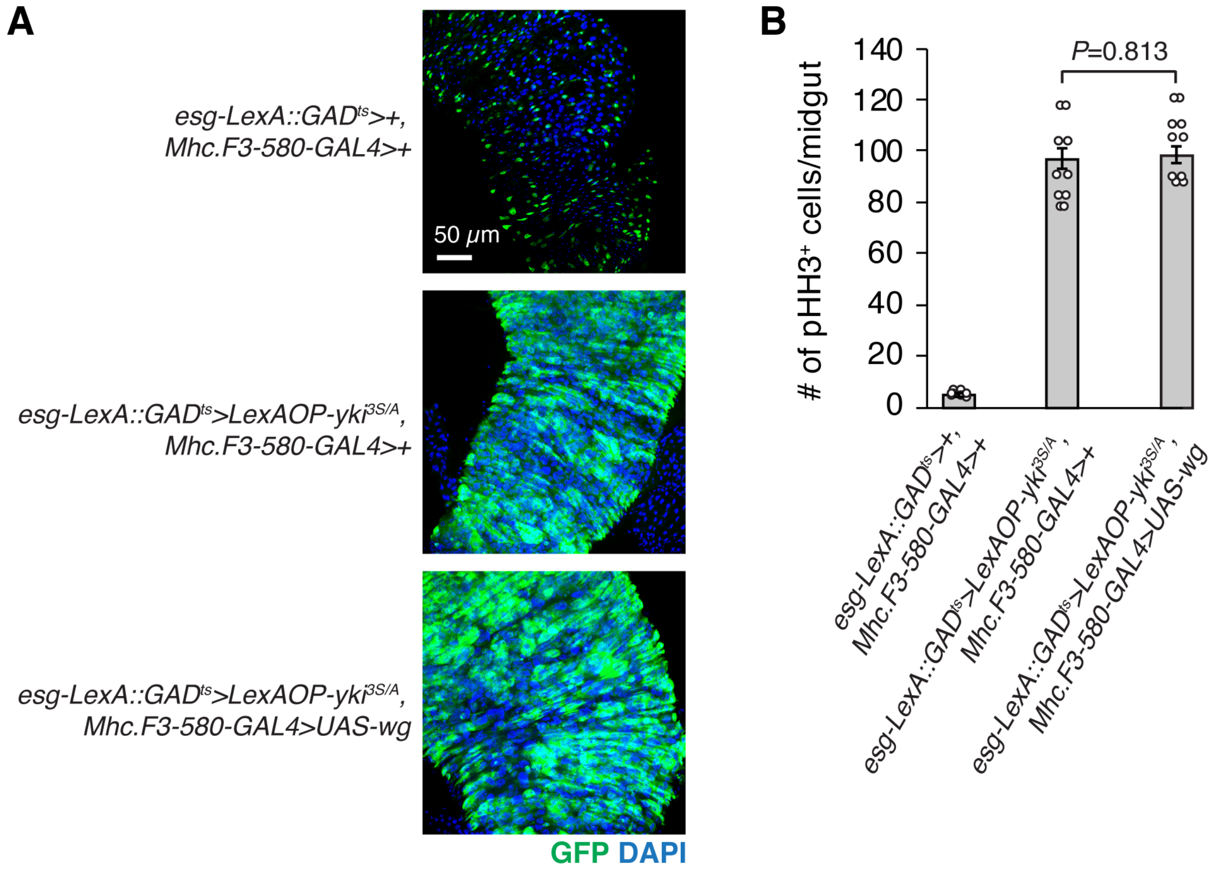


**Figure S4. Expression of Wg in the muscle has no effect on the growth of *yki*$^{3S/A}$ tumors in the midgut.**

(A) Representative images of posterior midguts. Transgenes were induced for 6 days. GFP (green) marks *esg*$^{+}$ cells, and nuclei are stained with DAPI (blue). Scale bar, 50 $\mu$m.

(B) Quantification of phospho-Histone H3$^{+}$ (pHH3$^{+}$) cells per midgut after 6 days of transgene expression. Mean±SEMs are shown. *P*-value is calculated by two-tailed unpaired Student's *t*-test between two groups indicated by bracket.

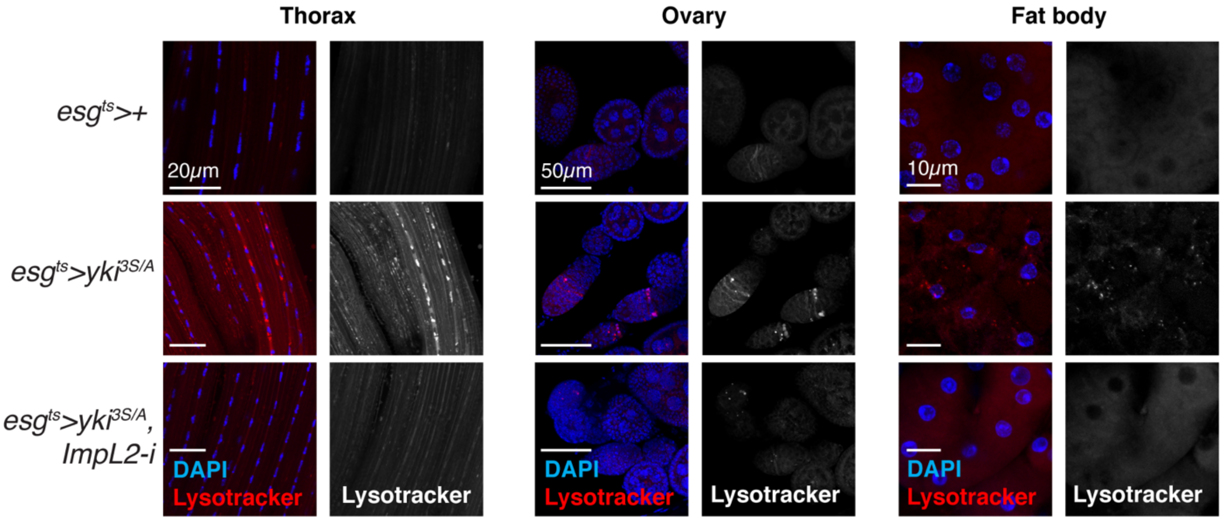


**Figure S5. Depletion of *ImpL2* in *yki*$^{3S/A}$ tumors is sufficient to suppress autophagy in the host tissues.**

Lysotracker staining in thorax, ovary, and fat body. Transgenes were expressed for 5 days. Tissues were stained with LysoTracker Red DND99 (red in merge), and counterstained for nuclei with DAPI (Blue). Scale bars are as indicated.
